# Coated Bacterial Vaccine Platform Overcomes Weak Antigen Immunogenicity: A Functional Approach to GnRH-Based Immunocastration

**DOI:** 10.64898/2026.04.20.719670

**Authors:** Inés Harguindeguy, Matías H Assandri, M. Antonieta Daza Millone, Sebastián F Cavalitto, María de los A Serradell, Gastón E. Ortiz

**Author notes:** **Corresponding author:** Prof. Dr. Gastón E. Ortiz or.

## Abstract

Immunocastration, a non-surgical strategy based on active immunization against gonadotropin-releasing hormone (GnRH), effectively suppresses steroidogenesis and spermatogenesis. However, peptide vaccines targeting poorly immunogenic antigens such as GnRH often fail to elicit robust adaptive immune responses, requiring adjuvants or carrier proteins. Previously, we introduced Coated Bacterial Vaccines (CBVs), a platform that uses chemically inactivated Gram-positive bacteria to display recombinant antigens fused to the SlpA carboxy-terminal domain (dSLPA) on their surface. This system leverages natural pathogen-associated molecular patterns (PAMPs) to enhance immunogenicity without additional adjuvants. In this work, we extended the application of the CBVs platform to enhance the immune response against a poorly immunogenic GnRH-based peptide vaccine. GnRH-CBVs were formulated using inactivated *Bacillus subtilis* var. *natto* coated with a recombinant GnRH tandem-repeat–dSLPA fusion protein and administered to male BALB/c mice. A chitosan-adjuvanted GnRH-dSLPA formulation served as a positive control. GnRH-CBVs induced a strong Th2-biased humoral response, characterized by predominant IgG1 levels comparable to those achieved with chitosan. The resulting antibodies effectively neutralized endogenous GnRH, reducing steroidogenesis and spermatogenesis and inducing marked testicular histological alterations. These findings support CBVs as a promising strategy to enhance peptide vaccine immunogenicity for veterinary immunocastration.

## 1. Introduction

Castration of male animals reared for meat production is a widespread practice in livestock farming to control aggressive and undesirable sexual behaviors [1,2]. In pig production, surgical castration is traditionally used to prevent boar taint, an unpleasant odor and taste of entire male pig meat mostly associated with the accumulation of androstenone and skatole in adipose tissue [3,4]. Although this conventional method is effective, it raises animal welfare concerns due to the pain, stress, and increased risk of postoperative infections, in addition to productivity losses [5,6]. These limitations have driven the development of less invasive, safer, and more ethical alternatives that ensure both animal welfare and meat quality [7].

In this context, immunocastration has emerged as a safe and effective non-surgical alternative to conventional castration [8]. This strategy involves inducing a specific immune response against gonadotropin-releasing hormone (GnRH), a hypothalamic decapeptide that stimulates the secretion of luteinizing hormone (LH) and follicle-stimulating hormone (FSH) from the anterior pituitary [9,10]. These hormones regulate the production of gonadal steroids, such as testosterone and androstenone, as well as spermatogenesis [11,12]. Active immunization against GnRH induces the production of neutralizing antibodies that block its interaction with pituitary receptors, interfering with the hypothalamic–pituitary–gonadal (HPG) axis. This leads to suppression of sexual steroid synthesis, gonadal development, and the accumulation of the compounds responsible for boar taint [13,14]. The effectiveness of this strategy has been validated in several animal species, including pigs, bulls, and companion animals [15–18].

Currently, several commercial immunocastration vaccines based on synthetic GnRH analogous are available, including Improvac and Bopriva® (Zoetis), Valora® (Ceva Santé Animale), and GonaCon™ (U.S. Department of Agriculture) [19–21]. However, the development of formulations capable of eliciting long-lasting autoimmune responses remains a challenge due to the low intrinsic immunogenicity of GnRH. A common approach has been the incorporation of highly immunogenic carrier proteins like diphtheria toxoid, tetanus toxoid, ovalbumin, or flagellin [22–24]. However, it has been described that the use of these carriers induces immunosuppression through an antigenic dominance effect, in which their immunodominant epitopes reduce the specific response against the target antigen [25,26]. This suppression could be overcome by replacing the carrier [27].

Another critical challenge in the development of peptide-based vaccines is the absence of pathogen-associated molecular patterns (PAMPs). These patterns, present in microbes, are essential for activating antigen-presenting cells and triggering an effective adaptive immune response [28]. To address this limitation, potent adjuvants are required not only to enhance antigen availability but also to generate the co-stimulatory signals necessary to activate the innate immune system and, subsequently, the lymphocyte response, thus inducing a robust and sustained humoral response [29–31]. In this regard, vaccines based on the presentation of antigens on the surface of bacterial carriers have emerged as a promising alternative to optimize antigen presentation and confer adjuvant properties to vaccine formulations. These vaccines naturally provide PAMPs and stimulate the activation of dendritic cells and macrophages [32,33].

Moreover, a key factor in achieving effective immunocastration is the immune profile induced. Previous studies have shown that a predominantly Th2 humoral response is required [11,34,35]. For these reasons, it is necessary to continue the design and development of new vaccine formulations, incorporating strategies that enhance antigen presentation to improve the efficacy of immunocastration vaccines.

On the other hand, a novel non-GMO antigen display platform called Coated Bacterial Vaccines (CBVs) has recently been developed. This system uses chemically inactivated Gram-positive bacteria to display recombinant antigens on their surface. This system enables efficient antigen presentation without the need for carrier proteins or adjuvants, enhancing the immunogenicity of peptide antigens through natural bacterial recognition mechanisms. In a previous study, tetanus toxin fragment C was used as a model antigen to validate this platform, showing that CBVs induce a Th2-polarized humoral response with high production of neutralizing antibodies and protective capacity against tetanus toxin challenge in mice [36].

Therefore, considering the potential of the CBVs platform to enhance immune responses and the limitations of current GnRH-based vaccines, this study aimed to evaluate the efficacy of CBVs in boosting the immune response against a weakly immunogenic GnRH peptide. To this end, a chemically inactivated *Bacillus subtilis* var. natto (ATCC 15245) carrier was coated with the GnRH tandem-repeat peptide, and the resulting GnRH-CBVs were evaluated in murine model. As a result, GnRH-CBVs elicited a Th2-polarized humoral response against the GnRH peptide, characterized by a predominance of IgG1 subclass antibodies. This immune response was associated with the suppression of steroidogenesis and spermatogenesis, as well as histological alterations in testicular tissue. These findings indicate that GnRH-CBVs can overcome the poor intrinsic immunogenicity of GnRH peptides and highlight their potential as a versatile platform for the development of safe and effective GnRH-based immunocastration vaccines.

## 2. Materials and methods

### 2.1. Antigen Design and Recombinant *E. coli* Generation

The GnRH-dSLPA chimeric antigen was constructed by fusing the carboxy-terminal cell wall-binding domain of the SlpA protein from Lactobacillus helveticus (dSLPA) to a tandem-repeat array of the GnRH decapeptide (QHWSYGLRPG). The repeats were connected by a (EAAAK)_4_ linker. The coding sequence was codon-optimized for *Escherichia coli* expression, chemically synthesized by General Biosystems, Inc. and cloned into the pET-29c(+) vector to obtain the expression plasmid pET-GnRH-dSLPA. Chemically competent *E. coli* BL21(DE3) cells were transformed with pET-GnRH-dSLPA by heat shock protocol, and positive transformants were selected on agar plates with Luria-Bertani broth (LB, pH 7.0) containing 50 µg/mL kanamycin, as described by Sambrook and Russel [37]. Colonies were screened using small-scale protein expression protocol [38]. The selected *E. coli* BL21 clone was cultured in LB-kanamycin medium and stored in 20% (v/v) glycerol solution at −80°C until use.

### 2.2. Antigen Production and Growth Conditions

For recombinant antigen production, *E. coli* BL21 cultures were grown in LB-kanamycin at 37°C, 200 rpm to an OD □□□□ of ∼0.6. Cultures were then cooled to 20°C, induced with 1 mM IPTG, and incubated for 8 h at 20°C, 200 rpm. Cells were harvested by centrifugation at 5,000 × *g* for 15 min at 4°C, resuspended to ∼40 OD □□□/mL in lysis buffer (50 mM Tris-HCl, pH 8.0; 150 mM NaCl; 2 mM EDTA; 1 mM PMSF; 0.5 mg/mL lysozyme; 1 μL/mL DNase), and incubated for 30 min at 37°C with agitation. Cells were disrupted using an ultrasonic homogenizer (SCIENTZ IID, model JY96-II, tip No. 6) at 30% amplitude for 30 cycles of 2 s ON / 2 s OFF. The homogenate was centrifuged at 32,000 × *g* for 30 min at 4°C to separate soluble (supernatant) and insoluble (pellet) fractions. The pellet was washed three times with PBS containing 2% Triton™ X-100 and 1 M urea, and inclusion bodies were solubilized in 8 M urea in PBS. Finally, 40 μL of each fraction was analyzed by SDS-PAGE.

### 2.3. Preparation of CBVs

CBVs were prepared as previously described by Harguindeguy et. al. [36]. Briefly, the bacterial carrier cell pellet obtained from 500 μL of 2×10□cells/mL of 2% (v/v) glutaraldehyde-inactivated *Bacillus subtilis* var. natto (ATCC 15245) was resuspended in 500 μL of the soluble GnRH-dSLPA fraction. The suspension was incubated at 80 rpm for 30 min at room temperature, and CBVs were recovered by centrifugation at 5,000 × *g* for 10 min at 4°C. Non-specifically bound proteins were removed by washing three times with 1 mL of PBS. The resulting CBVs were resuspended in 5 mL of PBS and stored at 4°C until further use. The ratio of GnRH-dSLPA bound to *B. subtilis* was quantified as protein amount per cell. Protein concentration was determined by SDS-PAGE followed by gel densitometry analysis, while cell concentration was quantified using a Neubauer chamber.

### 2.4. Purification of GnRH-dSLPA Antigen

GnRH-dSLPA was purified as previously described by Harguindeguy et. al. [36]. Briefly, the immobilized antigen was eluted from the bacterial carrier using 100 mM carbonate-bicarbonate buffer pH 10.2 for 15 min at 25°C. The eluate was dialyzed against PBS overnight at 4°C using a 12.4 kDa cut-off cellulose tubing and stored at -20°C until use. Protein concentration was determined using the Pierce BCA Protein Assay Kit (Thermo Fisher), and antigen integrity was assessed by SDS-PAGE.

### 2.5. Determination of Binding Kinetic Constants

Binding kinetics and affinity of the antigen-bacterial carrier interaction were analyzed by Multiparametric Surface Plasmon Resonance (MP-SPR) using a MP-SPR Navi^TM^ 210 (BioNavis) device. For this, the gold sensor surface (SPR102-Au) was treated with 200 µL of 0.01% (w/v) poly-L-Lysine, rinsed with Milli-Q water, and incubated with 100 µL of a 2 OD_600_/mL bacterial carrier suspension. After immobilization, the sensor was blocked with 500 µL of 5 mg/mL BSA, washed with Milli-Q water, and dried in a stream of nitrogen. All incubation steps were performed for 1 h at room temperature. Purified GnRH-dSLPA at increasing concentrations (2.5, 5, 10, 15, and 60 µg/mL in 0.05% (v/v) PBS-Tween 20) was injected at 10 µL/min for 10 min, and then washed for 10 min. Kinetic parameters were obtained from sensorgrams using TraceDrawer 1.5 software and a one-to-one interaction model. For each analyte concentration, the association and dissociation phases were independently analyzed, and the resulting fits were used to calculate the kinetic constants (*k*_a_, association rate and *k*_d_, dissociation rate) and the dissociation equilibrium constant (*K*_D_).

Additionally, the maximum binding capacity (*B*_max_) and apparent dissociation constant (*K*d) were determined by a saturation assay. A fixed number of bacterial carrier (1 × 10□cells) was incubated with increasing amounts of purified GnRH-dSLPA (12.5–125 μg in a final volume of 200 μL PBS) for 30 minutes at 25°C. After incubation, the cells were pelleted and washed thrice with PBS, and the amount of antigen bound to the bacterial surface was quantified as previously described by Harguindeguy et. al. [36].

### 2.6. Immunofluorescence Microscopy

Bacterial carrier and GnRH-CBVs were immobilized on coverslips pretreated with 0.01% (w/v) poly-L-Lysine and fixed with 4% (v/v) paraformaldehyde. Coverslips were incubated with 50 mM NH_4_Cl and blocked with 0.5% (w/v) BSA, and incubated with anti-GnRH-dSLPA mouse serum, followed by Alexa Fluor 488-conjugated goat anti-mouse IgG. Finally, samples were incubated with 1 µg/mL propidium iodide, and images were acquired using a confocal laser scanning microscope (TCS SP5; Leica). All incubation steps were performed for 30 min at room temperature using 200 μL of each solution, with three PBS washes between each step. The primary anti-GnRH-dSLPA serum was obtained from mice immunized with purified GnRH-dSLPA antigen following a standard immunization protocol [39].

### 2.7. Mice Immunization and Acute Toxicity Evaluation

Eight-week-old male BALB/c mice were used in two independent immunization trials. In both studies, animals were immunized intraperitoneally (i.p.) on days 0, 14, and 28 with 200 μL/dose of the respective vaccine. The first trial lasted 70 days and aimed to evaluate the effect of GnRH-CBVs on antibody production and gonadal function. The negative control group received *vaccine 1* (1 × 10^8^ cells/dose of bacterial carrier in PBS), the antigen control group received *vaccine 2* (20 μg/dose of GnRH-dSLPA in PBS), the positive control group received *vaccine 3* (20 μg/dose of GnRH-dSLPA in 0.5% (w/v) medium molecular weight chitosan in PBS), and the test group received *vaccine 4* (1 × 10^8^ GnRH-CBVs containing 20 μg/dose of GnRH-dSLPA in PBS). The second trial lasted 140 days and assessed the capacity of GnRH-CBVs to induce a long-lasting immune response. The negative control group received *vaccine 5* (2 × 10^8^ cells/dose of bacterial carrier in PBS), while the test group received *vaccine 6* (2 × 10^8^ GnRH-CBVs containing 40 μg/dose of GnRH-dSLPA in PBS). In both trials, blood samples were collected every 14 days and stored at -20°C.

Acute toxicity and GnRH-CBVs vaccine safety were evaluated by weekly monitoring of body weights during the second trial. In addition, mice were regularly observed following the initial inoculation to evaluate mortality, general condition, and behavioral changes [40].

### 2.8. Determination of IgG anti-GnRH Antibodies

An indirect ELISA was performed to measure total IgG, IgG1, and IgG2a in mouse serum. To prevent interference from anti-dSLPA antibodies, 2 μL of each serum sample were pre-incubated with 10 μL of dSLPA (0.4 mg/ml) for 1h at 25°C. Briefly, 96-well plates were coated with 500 ng/well of GnRH-dSLPA recombinant antigen or GnRH hormone (LHRH, Ilex Life Sciences) diluted in 100 mM carbonate-bicarbonate buffer pH 9.6 and incubated overnight at 4°C. After blocking with 200 μL/well of 3% (w/v) skim milk in PBS for 1 h at 30°C, plates were incubated with 50 μL/well of each serum diluted 1:500 (to detect antibodies against the recombinant antigen) or 1:100 (to detect antibodies against the endogenous hormone) in 1% (w/v) skim milk in PBS for 1 h at 30°C. Plates were then incubated with 50 μL/well of HRP-conjugated goat anti-mouse IgG, IgG1, or IgG2a antibodies diluted in 1% (w/v) skim milk in PBS for 1 h at 30°C. Finally, colorimetric reaction was developed with 50 μL/well of 2 mg/mL o-phenylenediamine substrate in 0.05 M citrate buffer pH 5 containing 0.03% (v/v) H_2_O_2_ for 10 min at room temperature, and stopped by adding 50 μL/well of 2 N H_2_SO_4_. Absorbance was measured at 492 nm using a Benchmark Plus microplate reader (Bio-Rad), and results were expressed as optical density (OD) values. Pre-immune sera absorbances were used as blanks. Between steps, plates were washed three times with 200 μL/well of PBS-Tween 20 and twice with 200 μL/well of PBS.

### 2.9. Measurement of Serum Testosterone Levels

To determine the concentration of testosterone in the serum of immunized mice, a chemiluminescent microparticle immunoassay (CMIA) was performed using the Architect 2nd Generation Testosterone assay (Abbott), according to the manufacturer’s instructions. The assay detection range was 0.04-19 ng/mL, with a lower limit of detection of 0.014 ng/mL and a limit of quantification of 0.023 ng/mL. Intra- and inter-assay coefficients of variation were 4.88% and 5.74%, respectively.

### 2.10. Testicular Morphological and Histological Analysis

At 70 (first trial) and 140 (second trial) dpi, animals were euthanized by CO_₂_ exposure, and testes were surgically extracted and fixed in 10% (v/v) buffered formalin. Testes were weighted in pairs using an analytical balance, and testicular volume was determined by liquid displacement. Briefly, each pair of testes was sequentially immersed in a 2-mL tube containing 1 mL of distilled water, and images were acquired before and after immersion while maintaining a fixed camera position and focus. Liquid column height was measured from the base of the tube to the lower meniscus using ImageJ software. The displaced volume was calculated as:

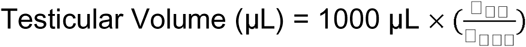

where h_wt_ and h_wot_ correspond to the liquid height with and without testes, respectively. Normalized testicular weight (NTW) and volume (NTV) were calculated as the ratios of testicular weight to body weight and testicular volume to body weight, respectively.

For histological analysis, testes were dehydrated, embedded in paraffin, and transversely sectioned into 5-µm-thick slices using a Leica RM2125 RTS microtome. Testicular tissue was stained with Hematoxylin–Eosin (H-E) and examined under a DMLB microscope coupled to a DC100 digital camera (Leica). Spermatid presence and seminiferous tubules diameter were evaluated in five representative histological sections from each testis. Quantitative image analysis was performed using ImageJ software.

### 2.11. Macrophage Culture and Stimulation Assays

RAW 264.7 murine macrophage (ATCC) were cultured in 24-well plates in Dulbecco’s modified Eagle’s medium (DMEM) supplemented with 10% (v/v) fetal bovine serum and 1% penicillin–streptomycin at 37°C in 5% CO_2_. For stimulation assays, cells (4 x 10^5^ cells/well) were incubated for 3, 6, or 24 h with either bacterial carrier (2 × 10^7^ cells/well), purified GnRH-dSLPA (4 µg/well), or GnRH-CBVs (2 × 10^7^ cells carrying 4 µg of GnRH-dSLPA/well). DMEM and 2 µg/mL lipopolysaccharide (LPS from *E. coli* O111:B4) were used as negative and positive controls, respectively. To minimize residual LPS effects, 10 μg/mL polymyxin B was added to all conditions except the positive control. Following treatment, cells were harvested by centrifugation at 3000g for 10 min, and both pellets and supernatants were collected for further analysis.

### 2.12. Quantitative Analysis of Interleukin mRNA Expression

Cell pellets collected at 3- and 6-h post-stimulation were used to evaluate cytokine mRNA expression by quantitative reverse transcription PCR (qRT-PCR). Total RNA was extracted using the RNAiso Plus Kit (Takara Bio), and cDNA was synthesized with the GoScript™ Reverse Transcriptase Kit (Promega) using 10 µM random primers, following the manufacturer’s instructions. The qRT-PCR was performed on CFX96 Touch Real-Time PCR (Bio Rad) using 5 µL of SYBR Green PCR Master Mix, 2 µL of cDNA (150 ng), and 0.6 μM of each primer (Table S1). Thermocycling conditions consisted of 94 °C for 5 min, followed by 39 cycles of 92 °C for 20 s and 60 °C for 20 s. Relative expression levels were calculated using the 2^−ΔΔCt^ method, with β-actin as the reference gene.

### 2.13. IL-6 Quantification

Interleukin-6 (IL-6) levels in macrophage culture supernatants were quantified after 24 h of stimulation using a commercial ELISA kit (BD Biosciences), according to the manufacturer’s instructions. The detection limit was 5 pg/mL.

### 2.14. Statistical Analysis

All results are reported as mean ± standard error of the mean (SEM), unless otherwise specified. Data analysis and graphing were performed using GraphPad Prism version 6.0 (GraphPad Software). Statistical significance between two groups was determined using Mann-Whitney *U* test, while multi-group comparisons data were analyzed by the one-way ANOVA followed by Dunnett’s or Tukey’s post-test. Differences were considered statistically significant at *p*<0.05.

## 3. Results

### 3.1. Expression of GnRH-dSLPA and Production of GnRH-CBVs

As shown in Figure 1A, the recombinant GnRH-dSLPA peptide was successfully expressed in *E. coli* BL21 after induction with 1 mM IPTG at 20°C. In comparison with the pre-induction sample, an overexpressed protein band of approximately 49 kDa corresponding to the molecular weight of recombinant GnRH-dSLPA peptide was detected in the post-induction sample. Moreover, it is possible to note that the recombinant peptide was present in soluble and insoluble fractions of the cell lysate. In addition, the induction conducted at 30°C or 37°C resulted in predominantly insoluble GnRH-dSLPA accumulation (data not shown).

**FIGURE 1.**
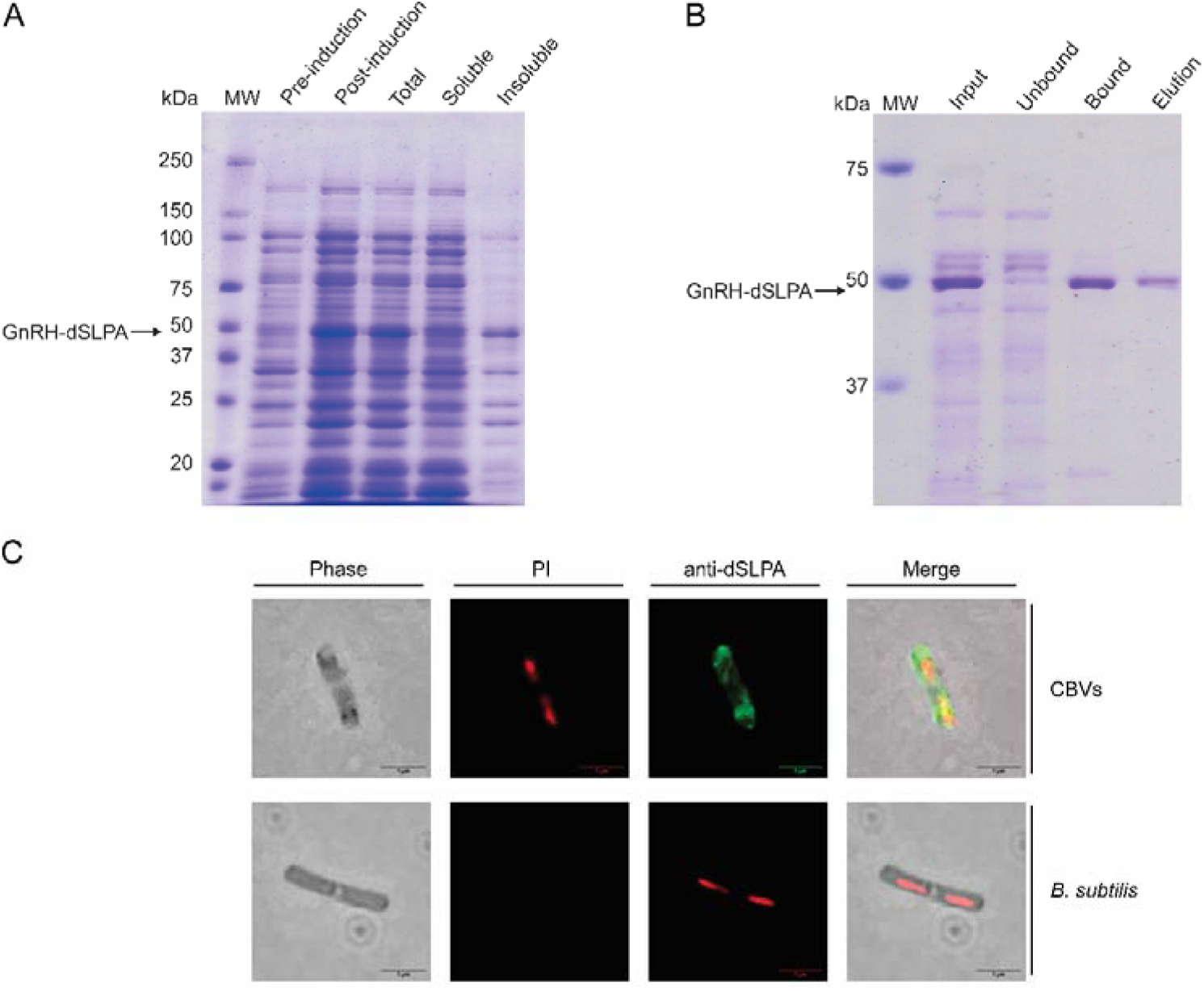
GnRH-CBVs production and characterization. (A) SDS-PAGE analysis of GnRH-dSLPA expression. Molecular weight marker (Precision Plus Protein BioRad, lane 1), total protein from *E. coli* lysate before (lane 2) and after induction (lane 3), total cell lysate (lane 4), soluble fraction (lane 5) and insoluble fraction (lane 6) after induction with 1 mM IPTG at 20°C for 8 h. (B) SDS-PAGE analysis of interaction. Molecular weight marker (Precision Plus Protein BioRad, lane 1), protein lysate input (lane 2), unbound protein fraction (lane 3), specific bound protein on *B. subtilis* surface (lane 4), eluted protein (lane 5). (C) Confocal immunofluorescence microscopy. CBVs and *B. subtilis* (negative control) stained with propidium iodide (red, PI panel), primary anti-dSLPA monoclonal and secondary anti-mouse Alexa Fluor 488 antibodies (green, anti-dSLPA panel). Scale bar: 1 µm.

Following cell lysis and fractionation, the soluble fraction was incubated with a bacterial carrier suspension to produce GnRH-CBVs. SDS-PAGE analysis revealed that the bacterial carrier efficiently retained GnRH-dSLPA from the *E. coli* lysate, as evidenced by the difference between the input and bound fractions (Figure 1B). This result demonstrates the specificity of the interaction between the antigen and the bacterial carrier.

To confirm this interaction, complementary confocal immunofluorescence microscopy. As shown in Figure 1C, a homogeneous fluorescent signal was observed across the entire bacterial carrier surface, suggesting a uniform antigen distribution and surface accessibility.

### 3.2. Evaluation of CBVs Antigen Binding Capacity

To further characterize the interaction between GnRH-dSLPA and the bacterial carrier a real-time binding analysis using MP-SPR was performed. The resulting sensorgrams exhibited well-defined association and dissociation phases (Figure 2A), confirming the ability of the antigen to associate with the immobilized bacterial carrier surface and enabling reliable kinetic analysis using a one-to-one interaction model. Individual curves fitting yielded an association rate constant (*k*_a_) of 7.4 ± 0.8 × 10³ M□¹ s□¹ and a dissociation rate constant (*k*_d_) of 3 ± 1 × 10□³ s□¹, corresponding to an equilibrium dissociation constant (*K*_D_) of 4 ± 1 × 10 □□M, which is consistent with a stable, high-affinity interaction under the tested conditions.

**FIGURE 2.**
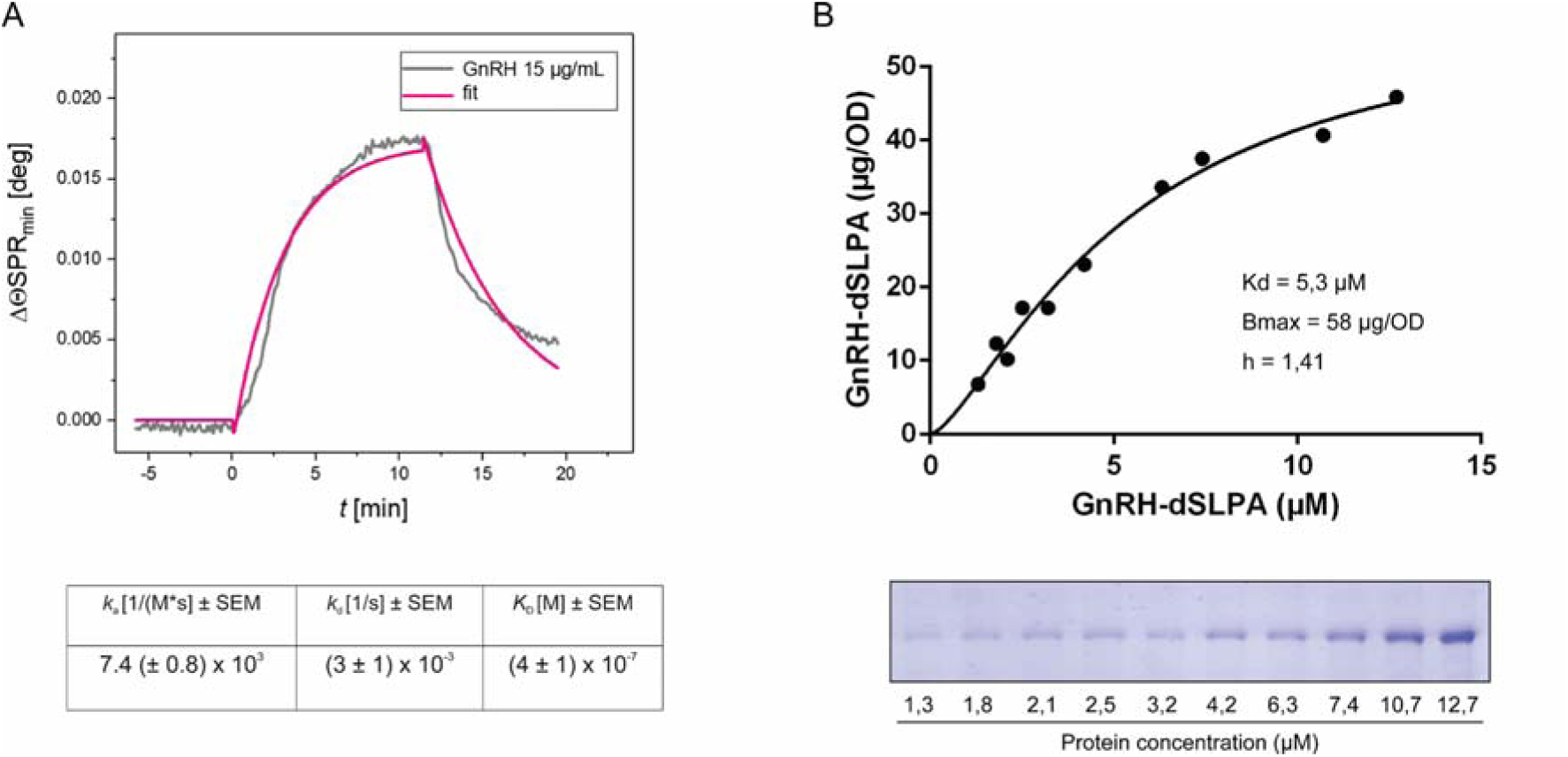
Binding kinetics parameters determination. (A) MP-SPR analysis. Representative sensorgram showing the change in SPR peak minimum angular position (ΔΘ_SPR min_) versus time during the interaction of GnRH-dSLPA (15 µg/mL) with immobilized bacterial carrier (gray line) and the corresponding global fit to a one-to-one interaction model (colored line). t=0 starts the injection of GnRH-dSLPA solution and t=10 returns to PBS-T buffer flow. Five analyte concentrations (2.5, 5, 10, 15, and 60 µg/mL) were measured under identical conditions, and *k*_a_, *k***_d_** and *K*_D_ were calculated from the averaged values obtained across the five independent fits (*n*=10). Mean ± SEM values are shown. (B) GnRH-dSLPA adsorption isotherm (up panel) was obtained using increasing concentrations of antigen (from 1.3 to 12.7 μM) and the bounded protein to bacteria was quantified by densitometry analysis of SDS-PAGE (down panel). The dissociation constant K_d_ (5.3 µM) and the maximum binding capacity B_max_ (58 µg/OD resin) were determined by nonlinear regression of the binding isotherm using GraphPad Prism 6.0 (GraphPad Software).

To further define antigen-loading capacity, an adsorption isotherm assay was performed by incubating a fixed amount of bacterial carrier cells with increasing concentrations of purified GnRH-dSLPA. Bound antigen was quantified by SDS-PAGE followed by densitometric analysis. Nonlinear regression of the resulting saturation curve, fitted to a Hill model, estimated an apparent dissociation constant (K_d_) of 5.30 μM, and a maximum binding capacity (B_max_) of 58 μg/OD, indicating a strong and specific interaction that allows extensive antigen immobilization on the bacterial carrier surface. As shown in Figure 2B, antigen association increased with GnRH-dSLPA concentration until it reached a plateau at approximately 10 μM. Although the apparent K_d_ obtained from the adsorption assay was one order of magnitude higher than the *K*_D_ determined by MP-SPR, both values fall within a comparable affinity range. Together, these results confirm the structural integrity and high antigen-binding capacity of the bacterial carrier, supporting CBVs suitability as a vaccine delivery platform.

### 3.3. Humoral Immune Response Induced by GnRH-CBVs

In the first immunization trial, we assessed whether GnRH-dSLPA formulated as CBVs could elicit an effective humoral response by measuring the level of serum antibodies against the recombinant antigen and the endogenous hormone over a 70-day period. As shown in Figure 3A, mice immunized with 20 µg/dose of GnRH-CBVs (*vaccine 4*) presented a significant increase and constant levels of specific IgG against the recombinant antigen, compared to those induced by either the bacterial carrier (*vaccine 1*) or 20 µg/dose of GnRH-dSLPA in PBS (*vaccine 2*). In contrast, animals immunized with *vaccine 2* exhibited only minimal antibody levels, while *vaccine 1* showed no detectable anti-GnRH-dSLPA IgG. In this regard, the antibody response profile elicited by GnRH-CBVs was comparable to that exhibited by GnRH–dSLPA chitosan formulation (positive control, *vaccine 3*).

**FIGURE 3.**
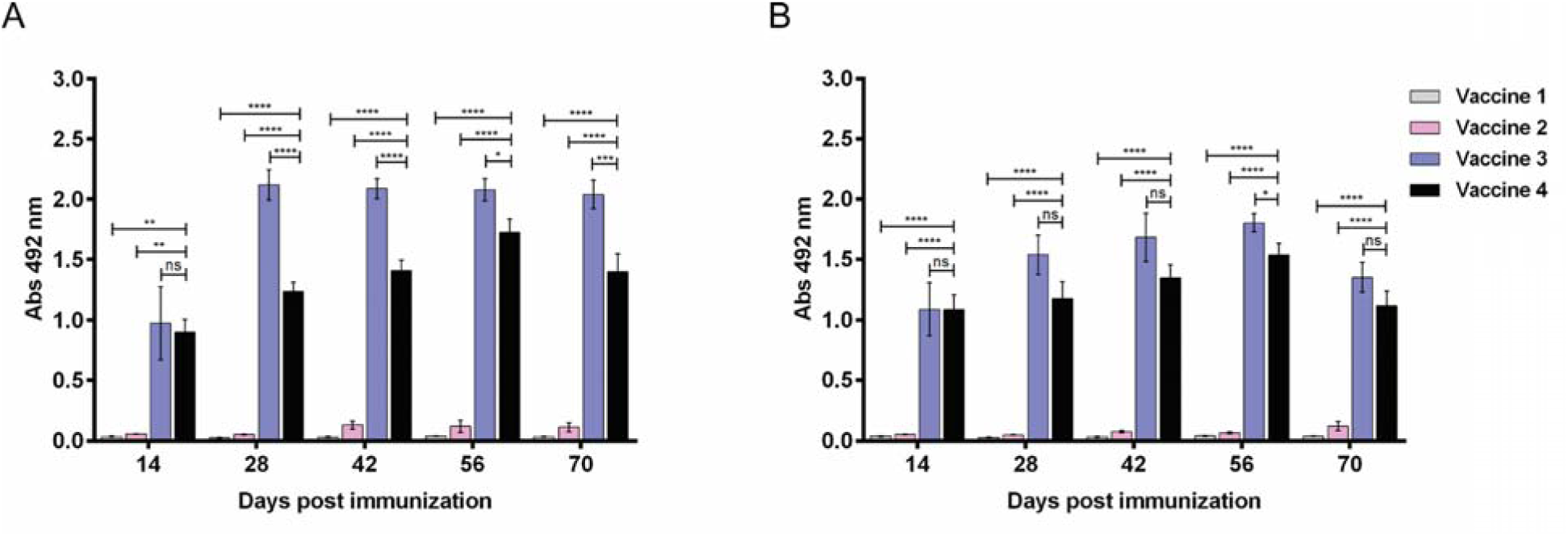
CBVs displaying GnRH-dSLPA induce a robust and sustained humoral immune response. Time course of serum IgG responses in BALB/c mice immunized with *vaccine 1* (1 × 10^8^ bacterial carrier cells), *vaccine 2* (20 _μ_g of GnRH-dSLPA), *vaccine 3* (20 _μ_g of GnRH-dSLPA in chitosan), or *vaccine 4* (1 × 10^8^ GnRH-CBVs containing 20 _μ_g of GnRH-dSLPA) on Days 0, 14 and 28. Serum samples were collected every 14 days until 70 dpi for specific antibodies anti-GnRH determination by ELISA. (A) IgG levels against the GnRH-dSLPA recombinant peptide. (B) IgG levels against native GnRH hormone. Antibody levels are expressed as absorbance values at 492 nm obtained from a 1:500 (A) or 1:100 (B) serum dilution. Data are shown as the mean ± SEM (*n*=5/group). Statistical significance was assessed using the one-way ANOVA followed by Tukey’s multiple comparison test (*p < 0.05, **p < 0.01, ***p < 0.001, ****p < 0.0001, ns; not significant).

On the other hand, as shown in Figure 3B a similar pattern was observed for anti-GnRH IgG against the native hormone, with antibody levels increasing after each booster immunization and remaining significantly higher than those of the negative control groups (*vaccine 1* and *vaccine 2*). Moreover, these responses were comparable to those elicited by positive control (*vaccine 3*).

These results suggest that GnRH-CBVs effectively enhance the humoral immune response against GnRH, achieving antibody levels comparable to those induced by a conventional GnRH–chitosan-adjuvanted vaccine. Furthermore, antibodies elicited by *vaccine 4* efficiently recognized native GnRH hormone, indicating their potential to function as neutralizing antibodies against GnRH.

### 3.4. Immune Response Profile Elicited by GnRH-CBVs

To characterize the immune profile induced by GnRH-CBVs, IgG1 and IgG2a levels against the recombinant antigen were quantified in sera collected on Day 70 in the first trial (Figure 4A). Mice immunized with GnRH-CBVs (*vaccine 4*) developed significantly higher IgG1 antibody levels compared to IgG2a. A similar isotype distribution was observed in the positive control (*vaccine 3*), whereas mice receiving GnRH in PBS (*vaccine 2*) exhibited only minimal antibody responses. The predominance of IgG1 over IgG2a suggests that immunization with GnRH-CBVs could promote a Th2-mediated immune response.

**FIGURE 4.**
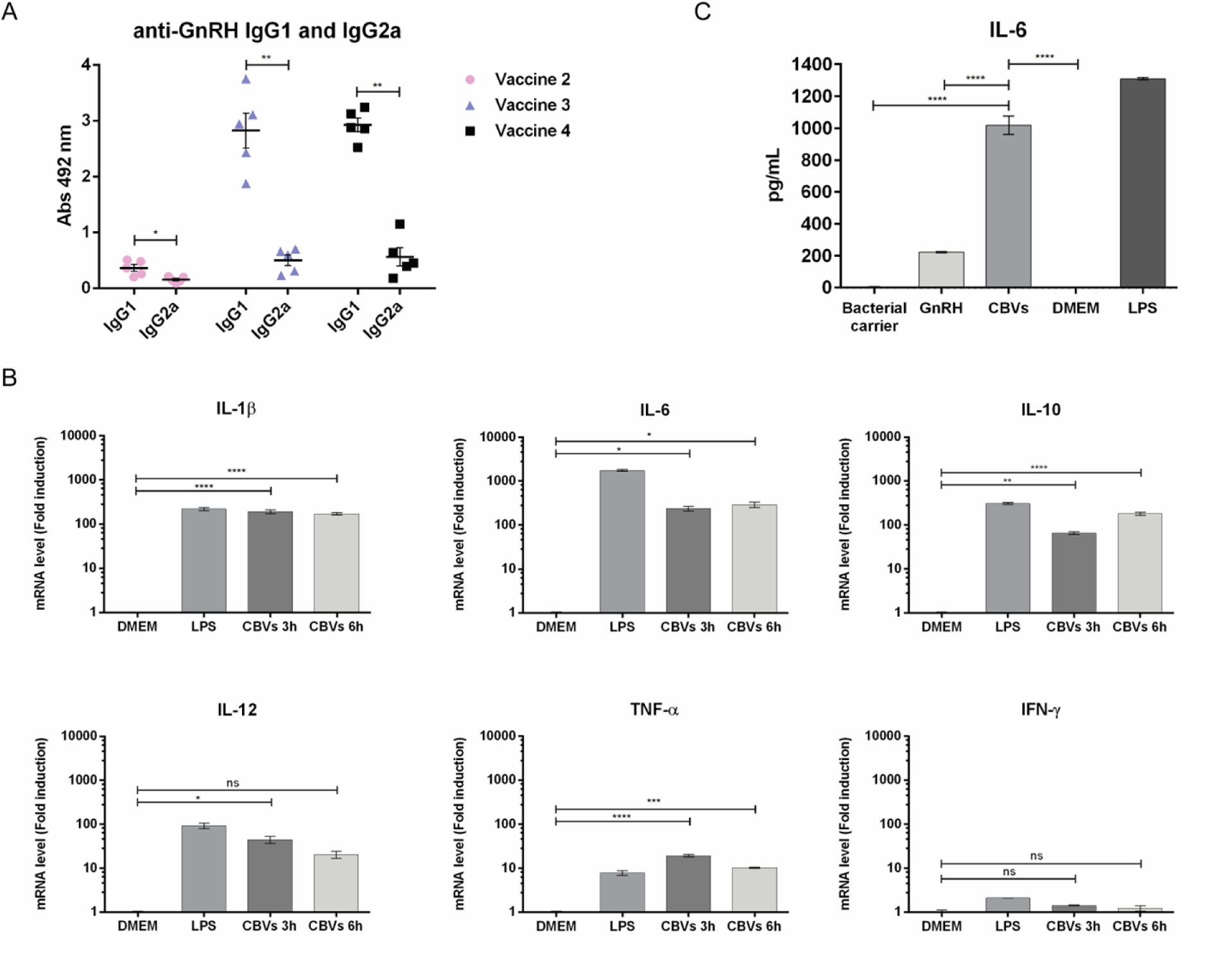
Immune polarization and macrophage activation induced by GnRH-CBVs. (A) Anti-GnRH IgG1 and IgG2a isotypes levels measured by ELISA at 70 dpi in first immunization trial, *vaccine 2* (GnRH in PBS), *vaccine 3* (GnRH in chitosan), or *vaccine 4* (GnRH-CBVs). Error bars indicate mean ± SEM of independent samples (*n*=5). Statistical significance was analyzed by Mann-Whitney *U* test (*p<0.05, **p<0.01). (B) Relative interleukin mRNAs expression levels determined by qRT-PCR at 3 and 6 h from cells stimulated with 2 × 10_GnRH-CBVs carrying 4 µg of GnRH-dSLPA, LPS (positive control), or DMEM (negative control). Data are presented as fold induction relative to unstimulated control (DMEM). Error bars indicate mean ± SEM of independent experiments (*n*=3). Statistical significance was assessed by one-way ANOVA followed by Dunnett’s multiple comparison test (*p < 0.05, **p < 0.01, ***p < 0.0001, ****p < 0.0001, ns; no significant). (C) Interleukin IL-6 production quantified by ELISA at 24 h from cells stimulated with 2 × 10_ bacterial carrier cells, 4 µg of GnRH-dSLPA, 2 × 10_ GnRH-CBVs carrying 4 µg of GnRH-dSLPA, LPS (positive control), or DMEM (negative control). Error bars indicate mean ± SEM of independent experiments (*n*=3). Statistical significance of CBVs with each treatment was assessed by one-way ANOVA followed by Tukey’s multiple comparison test (****p < 0.0001).

To further assess the direct activation of antigen-presenting cells by GnRH-CBVs and to characterize the early transcriptional changes in cytokine expression triggered by CBVs, mRNA expression levels of pro- and anti-inflammatory cytokines were quantified by RT-qPCR (Figure 4B). GnRH-CBVs stimulated RAW 264.7 macrophages, leading to a significant increase in IL-1β, IL-6, and IL-10 transcripts compared to unstimulated controls (170-, 290-, and 180-fold, respectively, at 6 h post-stimulation). In contrast, IL-12, TNF-α, and IFN-γ mRNA expression was induced to a lower extent (20-, 10-, and 1-fold, respectively, at 6 h post-stimulation).

Finally, cytokine secretion was evaluated by measuring IL-6 production 24 h after stimulation. As shown in Figure 4C, cells stimulated with GnRH-CBVs produced significantly increased IL-6 secretion at 24 h compared to cells treated with the bacterial carrier or purified GnRH-dSLPA.

In summary, GnRH-CBVs induced an IgG1-dominant antibody response *in vivo* and a cytokine mRNA profile in macrophages characterized by strong IL-1β, IL-6, and IL-10 induction with limited upregulation of IL-12, TNF-α, and IFN-γ. These profiles suggest a predominantly Th2-polarized humoral immune response. However, *in vivo* cytokine analysis is required to confirm these preliminary results.

### 3.5. Effect of GnRH-CBVs Immunization on Serum Testosterone Levels and Testicular Histomorphology

To determine the effect of GnRH-CBVs vaccination on gonadal function, serum testosterone levels and testicular histomorphology were analyzed in the different experimental groups at the end of the first immunization trial. Mice immunized with GnRH-CBVs (*vaccine 4*) exhibited a significant reduction in circulating testosterone levels compared with the groups immunized with bacterial carrier (*vaccine 1*) and GnRH-PBS (*vaccine 2*) (Figure 5A). As expected, the group that received GnRH-dSLPA chitosan formulation (positive control, *vaccine* 3) also induced a significant decrease in testosterone levels. It is important to note that no significant differences were observed between *vaccine* 4 and *vaccine 3*, suggesting comparable efficacy between both formulations in suppressing steroidogenesis (Figure 5A).

**FIGURE 5.**
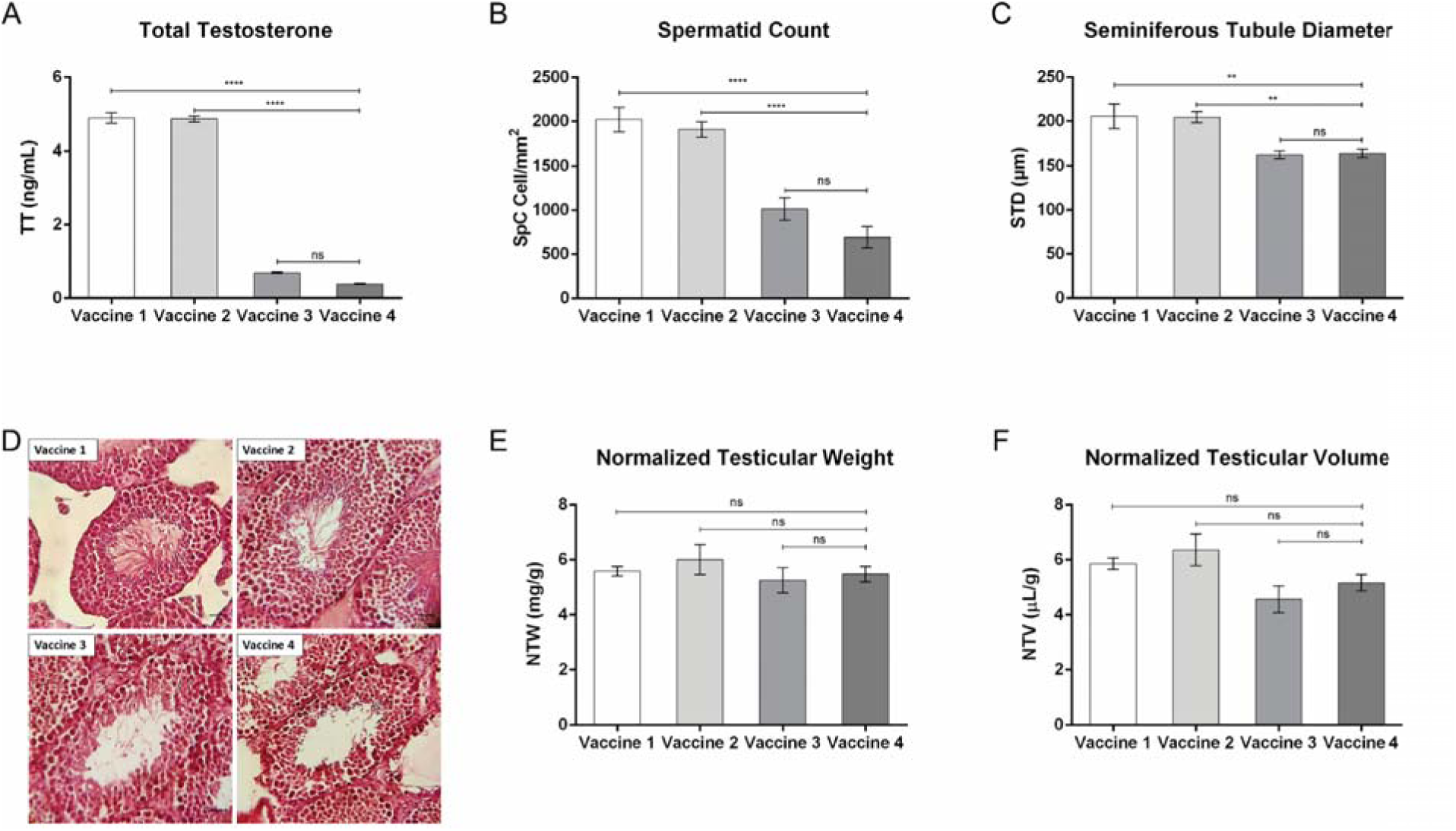
Analysis of serum testosterone levels, spermatogenesis, and testicular histo-morphology at 70 dpi. (A) Serum total testosterone concentrations (ng/mL), (B) spermatid counts per unit area (cell/mm^2^), and (C) seminiferous tubules diameter (µm) were quantified at the end of the first trial. (D) Representative H-E stained testicular sections are shown for each experimental group, illustrating preserved seminiferous tubule architecture in *vaccine 1* and *vaccine 2* and marked disruption of spermatogenesis in *vaccine* 3 and *vaccine* 4. Scale bar: 25 µm. (E) Normalized testicular weight (NTW, mg/g) and (F) normalized testicular volume (NTV, µL/g) were calculated relative to mice body weight. Data are expressed as the mean ± SEM. Statistical significance was determined using one-way ANOVA followed by Tukey’s multiple comparison test (**p < 0.01, ****p < 0.0001, ns, not significant).

The reduction in testosterone levels was associated with a hypogonadal phenotype, accompanied by alterations in testicular morphology of animals immunized with either *vaccine 4* or *vaccine 3*. In both groups, a significant decrease in spermatid counts and seminiferous tubules diameter was observed compared to mice vaccinated with *vaccine 1* or *vaccine 2* (Figure 5B and 5C). These results suggest a functional gonadal suppression consistent with the reduction of testosterone level and GnRH neutralization.

Histological examination further revealed severe degeneration of seminiferous tubules in the GnRH-CBVs (*vaccine 4*) and GnRH-chitosan (*vaccine 3*) groups, characterized by reduced tubular diameter, epithelial thinning with loss of the organized stratified germinal epithelium, enlargement of tubular lumens, and depletion of spermatogenic cells (Figure 5D). In contrast, testes from animals receiving the bacterial carrier (*vaccine 1*) or GnRH-PBS (*vaccine 2*) displayed well-organized seminiferous tubules architecture, intact germinal epithelium, and abundant spermatogenic cells (Figure 5D).

No statistically significant differences were detected among groups regarding macroscopic morphological changes associated with testicular atrophy, including normalized testicular weight and volume. Nevertheless, a trend toward reduced testicular weight and volume was observed in *vaccine 3* and *vaccine 4* groups, which may indicate an early stage of the atrophic process (Figure 5E and 5F). Overall, the endocrine and morphological results demonstrate that GnRH-CBVs formulation induces a marked immunocastration effect, characterized by reduction in testosterone levels, suppression of spermatogenesis, and structural alterations of the seminiferous tubule. The strong correlation between histo-morphological alterations and endocrine suppression underscores the efficacy of the CBV-based delivery platform in efficiently impairing gonadal function.

### 3.6. Long-Term Immunogenicity and Safety of GnRH-CBVs

To evaluate the durability of the immune response induced by the CBV platform, a second immunization trial was conducted using a higher dose GnRH-CBVs formulation. As shown in Figure 6A, mice immunized with 40 µg/dose of GnRH-CBVs (*vaccine 6*) developed a robust response against the recombinant antigen that was detectable after the first immunization and increased progressively following booster administrations. In contrast, no anti-GnRH-dSLPA IgG antibody response was detected in animals receiving the bacterial carrier alone (*vaccine 5*). Similarly, anti-GnRH IgG levels followed a comparable kinetic profile, with a progressive increase after the booster vaccination and sustained antibody response up to day 140 (Figure 6B). Although a moderate decline was observed at later time points, antibody levels remained markedly above the control values. These results indicate that antigen presentation through the CBV platform induces a durable and sustained humoral immune response.

**FIGURE 6.**
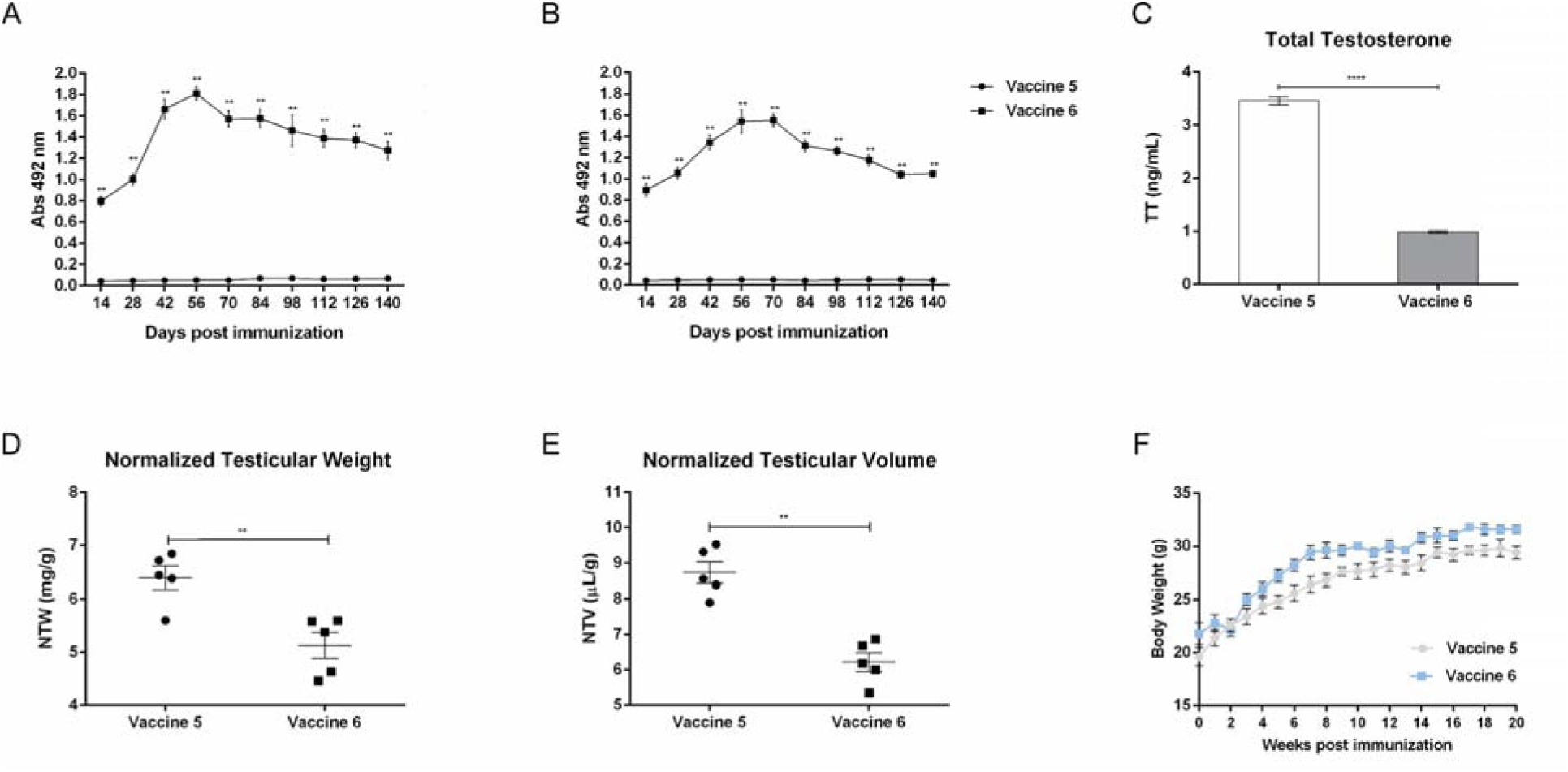
Second immunization trial for the evaluation of long-term immunogenicity and functional effects of GnRH-CBVs. Mice were immunized with *vaccine 5* (2 × 10^8^ bacterial carrier cells) or *vaccine 6* (2 × 10^8^ GnRH-CBVs containing 40 μg/dose of GnRH-dSLPA) on Days 0, 14 and 28. Serum samples were collected every 14 days until 140 dpi for specific antibodies anti-GnRH determination by ELISA. At 140 dpi, mice were sacrificed, serum testosterone levels were measured, and testes were collected for morphometric analysis. (A) IgG levels against the GnRH-dSLPA recombinant peptide. (B) IgG levels against native GnRH hormone. Antibody levels are expressed as absorbance values at 492 nm obtained from a 1:500 (A) or 1:100 (B) serum dilution. (C) Serum total testosterone concentrations (ng/mL) at the end of the second trial. (D) Normalized testicular weight (NTW, mg/g) and (E) normalized testicular volume (NTV, μL/g) were calculated relative to mice body weight. (F) Negative control (*vaccine 5*) and GnRH-CBVs (*vaccine 6*) immunized mice were weighed every 14 days post first immunization until 140 dpi. All data are expressed as the mean ± SEM (*n*=5/group). Mann-Whitney U Test was performed to analyze the statistical significance of results (*p < 0.05, **p < 0.01, ****p < 0.0001).

Consistent with the sustained humoral response, mice immunized with GnRH-CBVs exhibited a significant reduction in circulating testosterone levels compared to the negative control (Figure 6C), indicating long-term functional suppression of gonadal activity. Morphometric analysis revealed that the normalized testicular weight (Figure 6D) and volume (Figure 6E) were significantly lower in the GnRH-CBVs group than in the bacterial carrier group.

Furthermore, to assess acute toxicity of GnRH-CBVs vaccine, changes in live body weight and general health conditions were monitored during the second trial. Briefly, the animals showed no adverse reactions at the inoculation sites or behavioral alterations, and body weight progression followed a normal growth trend in both groups, with no statistically significant differences between them (Figure 6F). Taken together, these results suggest that immunization with GnRH-CBVs is safe and can induce a long-lasting humoral immune response that persists for at least 140 days and is associated with sustained endocrine and gonadal suppression.

## 4. Discussion

A major challenge in the development of effective immunocastration vaccines is overcoming the low immunogenicity of self-antigens such as GnRH. As a small peptide, GnRH vaccine formulations often require strong adjuvants or carrier proteins to induce neutralizing antibody responses [41]. Over the past years, several strategies have been explored to address this limitation, including conjugation to carrier proteins like CRM197 [42], keyhole limpet hemocyanin (KLH) [23], or mycobacterial heat shock protein 70 [43], often formulated with oil-based adjuvants [10] or water-in-oil-in-water emulsions. However, these approaches often suffer from antigenic dominance, where the carrier protein suppresses the specific response against the target peptide. More recently, bacterial-based delivery systems have emerged as promising alternatives. For instance, *Salmonella* flagellin-conjugated GnRH constructs induced strong anti-GnRH antibody responses and marked testicular suppression in pre-pubertal boars [44], while bacteriophage T7 vectors elicited antibodies that reduced testosterone levels and inhibited spermatogenesis in murine model [45]. Likewise, oral attenuated Salmonella vectors have demonstrated immunogenicity, inducing anti-GnRH antibodies and testicular atrophy in rodents [46].

In a previous study, we introduced the Coated Bacterial Vaccines (CBVs) platform and demonstrated its ability to induce a strong immune response using tetanus toxin fragment C (TTFC) as a model antigen [36]. Based on these findings, in this work we continue exploring this antigen delivery system by evaluating its capacity to elicit an effective immune response against a poorly immunogenic GnRH-based antigen and to assess its functional efficacy in a murine model of immunocastration.

To this end, the GnRH-dSLPA chimeric antigen was successfully recovered from the soluble fraction of *E. coli* lysates, in contrast to previous reports describing the accumulation of GnRH-based fusion proteins in insoluble inclusion bodies when expressed in bacterial systems [47]. This favorable expression profile may be attributed to codon optimization, low-temperature induction conditions, and the intrinsic solubility-enhancing properties of the dSLPA domain. These factors likely contribute to improved downstream processing while preserving the native conformation required for high-affinity binding to the bacterial carrier surface.

The interaction between the dSLPA domain and glutaraldehyde-inactivated *B. subtilis* surface proved to be stable and specific, with an equilibrium dissociation constant (*K*_D_) in the submicromolar range (4 ± 1 × 10⁻L M) as determined by SPR analysis. This high-affinity interaction is consistent with our previous validation of the CBV platform using TTFC-dSLPA antigen, where a similar kinetic parameter (K_d_ = 4.70 μM) [36]. These results confirm that the dSLPA domain consistently directs a reproducible and homogeneous display of diverse antigens, a property crucial for ensuring consistent and predictable immune recognition. Furthermore, the relatively high binding capacity (58 μg/OD) demonstrates that the CBV platform enables high-density antigen loading, which is critical for optimal immune presentation.

In this sense, our findings demonstrate that GnRH-CBVs induced a strong and sustained antibody response against both the recombinant GnRH-dSLPA fusion peptide and the native GnRH hormone. In contrast, mice immunized with soluble GnRH-dSLPA in PBS failed to develop detectable antibodies, underscoring the poor intrinsic immunogenicity of this antigen and highlighting the important role of the delivery system. Notably, the antibody levels elicited by GnRH-CBVs were comparable to those induced by the chitosan-adjuvanted GnRH formulation [11]. The ability of vaccine-induced antibodies to recognize native GnRH is critical for functional immunocastration, as it indicates that the antibodies can neutralize endogenous GnRH and disrupt the HPG axis. These results demonstrate that the CBVs platform can be as effective as conventional adjuvants, leveraging the natural PAMP-rich bacterial surface to provide the necessary co-stimulatory signals for immune activation [28,33].

The analysis of antibody IgG isotype distribution revealed a predominance of IgG1 over IgG2a, suggesting a Th2-polarized immune response. This polarization is consistent with previous reports suggesting that Th2 dominated responses are more effective for immunocastration, as they favor high-titer, high-affinity antibody production against GnRH [34,35]. This Th2 immune profile observed with GnRH-CBVs aligns with our earlier findings using the TTFC model antigen, where immunization with TTFC-CBVs also led to a significant predominance of IgG1 over IgG2a antibodies [36]. The fact that this Th2-polarized response is consistently elicited by two such distinct antigens, a strongly immunogenic toxin fragment (TTFC) and a weakly immunogenic self-peptide (GnRH), suggests that the CBV platform itself is the primary driver of this polarization. The innate immune signals provided by the *B. subtilis* carrier, such as its surface teichoic acids and exopolysaccharides, likely play a decisive role in shaping this response [48–50].

This Th2 response was further supported by *in vitro* macrophage stimulation, where GnRH-CBVs induced a significant upregulation of IL-1β, IL-6, and IL-10 mRNA, with only modest increases in IL-12, TNF-α, and IFN-γ. This profile is remarkably similar to the cytokine profile elicited by TTFC-CBVs in our previous study, reinforcing the concept that the CBV platform consistently instructs antigen-presenting cells towards a phenotype that favors a humoral, Th2-type adaptive immune response [36]. In addition, this immune profile is consistent with those reported in previous studies describing effective immunocastration [35]. It has been demonstrated that the immune profile induced by GnRH vaccination is crucial for determining effects on gonadal function and fertility, with Th2 responses associated with sustained antibody production and effective immunocastration [34]. However, *in vivo* cytokine analysis or T-cell phenotyping would provide more definitive evidence of the immune polarization.

The Th2 immune response observed is consistent with those reported in previous studies of effective immunocastration, evidenced by the suppression of gonadal activity [34,35]. In our experiments, the mice immunized with GnRH-CBVs exhibited significantly reduced serum testosterone levels compared to controls, alongside marked histological alterations in testicular tissue, including decreased spermatid counts, reduced seminiferous tubule diameter, and disruption of germinal epithelium architecture. These changes are consistent with GnRH neutralization and subsequent failure of the HPG axis [10,13]. Importantly, the magnitude of gonadal suppression was similar to that achieved with the chitosan adjuvanted formulation, confirming that CBVs are at least as effective as a standard adjuvant.

Finally, the long-term immunization trial further demonstrated the durability of the immune response, with anti-GnRH antibodies remaining detectable for at least 140 days post-immunization. These sustained antibody levels correlated with persistent testosterone suppression and testicular atrophy, confirming that this platform can maintain an effective hypogonadal state over a period relevant for practical applications in livestock, where a single vaccination course should ideally cover the production cycle [7,8]. An additional advantage of the CBV platform over conventional carrier protein-based vaccines is its ability to avoid carrier-induced epitopic suppression, which further supports the long-lasting immunity observed. It has been well documented that strong immune responses against carrier proteins such as tetanus toxoid or diphtheria toxoid can dominate the response and dampen antibody production against the linked weak antigen [25,26]. By displaying the antigen directly on the bacterial surface without a highly immunogenic protein carrier, CBVs circumvent this problem, thereby focusing the immune response on the target GnRH peptide and promoting long-term immunity. Our results, showing strong and sustained anti-GnRH responses without evidence of suppression, together with the absence of adverse effects on body weight or general behavior, support this hypothesis.

Despite these promising results, this study was conducted in a murine model, and extrapolation to target species such as pigs, cattle, cats or dogs will require further validation.

## 5. Conclusion

In this study we have demonstrated that the CBVs platform is an effective and adaptable strategy for enhancing the immunogenicity of weakly immunogenic antigens, as demonstrated with a GnRH-based immunocastration vaccine in a murine model. The GnRH-CBVs formulation elicited a robust and sustained humoral response, comparable to that achieved with a conventional adjuvant, which effectively suppressed gonadal function including reduced testosterone levels, histological alterations in testicular tissue, and suppression of spermatogenesis. The response was characterized by a predominance of IgG1 over IgG2a antibodies, indicative of a Th2-polarized profile consistent with effective immunocastration. The platform offers several benefits, including high-density antigen display, natural adjuvanticity through bacterial PAMPs, and the potential to avoid carrier-induced epitopic suppression. The long-lasting immune response and safe profile highlight the potential of CBVs in veterinary immunocastration. These findings demonstrate that the CBV platform effectively overcomes the poor intrinsic immunogenicity of GnRH. Beyond this application, the CBV platform represents a promising approach for the development of peptide-based vaccines targeting other weakly immunogenic antigens. Future research should focus on evaluating the efficacy of this technology in target species and exploring alternative routes of administration to optimize its practical utility.

## Authors Contributions

IH performed most of the experimental work, generated the figures, and significantly contributed to the writing and discussion of the manuscript. MHA conducted the macrophage activation experiments. MAS and SFC participated in the writing. GOE, as the principal investigator and project director, contributed to the experimental design, interpretation of the results, and writing of the manuscript. All authors contributed to the development of the article and approved the final submitted version.

## Ethics Statement

The experimental procedures of this study (protocol 010-42-24) were approved by the Institutional Animal Care and Use Committee of Exact Sciences Department at the National University of La Plata. Mice were immunized and challenged under the recommendations for animal experimentation (Helsinki Declaration and its amendments, and National Institutes of Health, United States: Guide for the Care and Use of Laboratory Animals).

## Abbreviations

The following abbreviations are used in this manuscript:

CBVs: Coated Bacterial Vaccines
dSLPA: carboxy-terminal fragment of SlpA protein
GnRH: Gonadotropin-Releasing Hormone
GnRH-dSLPA: GnRH-tandem-repeat peptide fused to dSLPA
HPG: Hypothalamic–Pituitary–Gonadal
TTFC: Tetanus Toxin Fragment C
TTFC-dSLPA: Tetanus Toxin Fragment C fused to dSLPA

## Data Availability Statement

The data that supports the findings of this study are available in the supplementary material of this article.

