## Supplemental Table 1 for "Coated Bacterial Vaccine Platform Overcomes Weak Antigen Immunogenicity: A Functional Approach to GnRH-Based Immunocastration"

**Supplementary**

**Table S1.** Sequence of primers used in real-time PCR.

| Gene | Protein name | Sense | Antisense |
| --- | --- | --- | --- |
| IL-1β | Interleukin 1β | GAAATGCCACCTTTTGACAGTG | TGGATGCTCTCATCAGGACAG |
| IL-6 | Interleukin 6 | GTTCTCTGGGAAATCGTGGAAA | AAGTGCATCATCGTTGTTCATACA |
| IL-10 | Interleukin 10 | CATTTGAATTCCCTGGGTGAGA | TGCTCCACTGCCTTGCTCTT |
| IL-12 | Interleukin 12 | GCAAAGAAACATGGACTTGAAGTTC | CACATGTCACTGCCCGAGAGT |
| TNF-α | Tumor necrosis factor-α | CATCTTCTCAAAATTCGAGTGACAA | CCTCCACTTGGTGGTTTGCT |
| IFN-Ɣ | Interferon-Ɣ | TGGCATAGATGTGGAAGAAAAGAG | TGCAGGATTTTCATGTCACCAT |
| Actin | β-actin | CGTCATCCATGGCGAACTG | GCTTCTTTGCAGCTCCTTCGT |
